# A novel, lightweight drive implant for chronic tetrode recordings in juvenile mice

**DOI:** 10.1101/2023.01.04.522760

**Authors:** Robert J Pendry, Lilyana D Quigley, Lenora J Volk, Brad E Pfeiffer

**Affiliations:** Department of Neuroscience, UT Southwestern Medical Center, Dallas, TX; Neuroscience Graduate Program, UT Southwestern Medical Center, Dallas, TX; O’Donnell Brain Institute, UT Southwestern Medical Center, Dallas, TX; Department of Psychiatry, UT Southwestern Medical Center, Dallas, TX

**Keywords:** *In vivo* physiology, Tetrode, Juvenile, Mice, LFP, Single Units, Micro-drive, Implant, Surgery, Recovery

## Abstract

**SHORT ABSTRACT:** We describe a novel micro-drive design, surgical implantation procedure, and post-surgery recovery strategy that allows for chronic field and single-unit recordings from up to sixteen brain regions simultaneously in juvenile and adolescent mice across a critical developmental window from p20 to p60 and beyond.

**LONG ABSTRACT:** *In vivo* electrophysiology provides unparalleled insight into sub-second-level circuit dynamics of the intact brain and represents a method of particular importance for studying mouse models of human neuro-psychiatric disorders. However, such methods often require large cranial implants which cannot be used in mice at early developmental timepoints. As such, virtually no studies of *in vivo* physiology have been performed in freely behaving infant or juvenile mice, despite the fact that a better understanding of neurological development in this critical window is likely to provide unique insights into age-dependent developmental disorders such as autism or schizophrenia. Here, we describe a novel micro-drive design, surgical implantation procedure, and post-surgery recovery strategy that allows for chronic field and single-unit recordings from up to sixteen brain regions simultaneously in mice as they age from postnatal day 20 (p20) to postnatal day 60 (p60) and beyond, a time window roughly corresponding to human ages 2-years-old through adult. The number of recording electrodes and final recording sites can be easily modified and expanded, allowing flexible experimental control of *in vivo* monitoring of behavior- or disease-relevant brain regions across development.

## INTRODUCTION

The brain undergoes large-scale changes during the critical developmental windows of childhood and adolescence^1–3^. Many neurologic and psychiatric diseases, including autism and schizophrenia, first manifest behaviorally and biologically during this period of juvenile and adolescent brain development^4–6^. While much is known regarding the cellular, synaptic, and genetic changes that occur across early development, comparatively little is known regarding how circuit- or network-level processes change throughout this time window. Importantly, circuit-level brain function, which ultimately underlies complex behaviors, memory, and cognition, is a non-predictable, emergent property of cellular and synaptic function^7–10^. Thus, to fully understand network-level brain function, it is necessary to directly study neural activity at the level of an intact neural circuit. In addition, to identify how brain activity is altered throughout the progression of neuropsychiatric disorders, it is critical to examine network activity in a valid disease model during the specific temporal window when behavioral phenotypes of the disease manifest and track observed changes as they persist into adulthood.

One of the most common and powerful scientific model organisms is the mouse, with large numbers of unique genetic strains that model neurodevelopmental disorders with age-dependent onset of behavioral and/or mnemonic phenotypes^11–21^. While it is challenging to correlate precise developmental time points between the brains of humans and mice, morphological and behavioral comparisons indicate that p20-21 mice represent human ages 2 to 3 years old and p25-p35 mice represent human ages 14-11, with mice likely reaching the developmental equivalent of a human 20-year-old adult by p60^3, 22^. Thus, to better understand how the juvenile brain develops and to identify how the neural networks of the brain become dysfunctional in diseases like autism or schizophrenia, it would be ideal to directly monitor brain activity *in vivo* in mice across the ages of 20 to 60 days old.

However, a fundamental challenge to monitoring brain activity across early development in mice is the small size and relative weakness of juvenile mice. Chronic implantation of electrodes, which is necessary for longitudinal studies of brain development, typically requires large, bulky housing to protect the fine electrode wires and interface boards^23, 24^, and the implants must be firmly attached to the mouse skull, which is thinner and less rigid in young mice due to reduced ossification. Thus, virtually all studies of *in vivo* rodent physiology have been performed in adult subjects due to their relative size, strength, and skull thickness. To date, most studies exploring *in vivo* juvenile rodent brain physiology have been performed in wild-type juvenile rats, which necessarily limits the ability to experimentally monitor juvenile brain function in a freely behaving model of a human disorder^25–30^.

In this manuscript, we describe novel implant housing, surgical implantation procedure, and post-surgery recovery strategy to chronically study long-term (up to four or more weeks) *in vivo* brain function of juvenile mice across a developmentally critical time window (p20 to p60 and beyond). Our implantation procedure allows for reliable, permanent affixation to the skull of juvenile mice. Furthermore, our novel micro-drive design is lightweight, and with minimal counter-balancing to offset the weight of the implant, does not impact behavioral performance of juvenile mice on typical behavioral paradigms.

## PROTOCOL

### 1. Micro-Drive Design and Construction

1.1) Digitally design and print the micro-drive (**Figure 1**)
  1.1.1) Download the micro-drive model templates from https://github.com/Brad-E-Pfeiffer/JuvenileMouseMicroDrive/releases/tag/v1.0.1.
  1.1.2) Identify the stereotaxic locations of target brain region(s) in an appropriate stereotaxic atlas.
  1.1.3) Using three-dimensional computer-aided design (3D CAD) software, load the template micro-drive cannula (**Figure 1B**).
  1.1.4) If necessary, modify the output cannula output locations on the micro-drive cannula model to target desired brain region(s). Each cannula hole extrusion should be at least 2 mm long to ensure that the tetrode will emerge from the cannula hole aiming straight at the target. The micro-drive cannula template is designed to bilaterally target the anterior cingulate cortex (one tetrode per hemisphere), hippocampal area CA1 (4 tetrodes per hemisphere), and hippocampal area CA3 (2 tetrodes per hemisphere), with one reference tetrode per hemisphere positioned in the white matter above hippocampal area CA1.
  1.1.5) If necessary, modify the micro-drive body (**Figure 1A**) to accommodate attachment of the electronic interface board (EIB). The micro-drive body template is designed to be used with a Neuralynx EIB model EIB-36-16TT or EIB-36-Narrow.
  1.1.6) Print the micro-drive body, cannula, cone, and lid at high resolution on a 3D printer (ideally with resolution better than 25 μm) and prepare the printed materials according to manufacturer protocols. Use printer resins with high rigidity.
1.2) Assemble the custom screws and attachments (**Figure 2A-B**)
  1.2.1) Using 3D CAD software, load the screw attachment models (**Figure 1E**).
  1.2.2) Print the screw attachments at high resolution on a 3D printer (ideally with resolution at least 25 μm) and prepare the printed materials according to manufacturer protocols. Use printer resins with high rigidity.
  1.2.3) Affix the screw attachments to each tetrode-advancing screw (**Figure 1F**) (Tetrode-advancing screws are custom manufactured at a machine shop prior to micro-drive construction.) Two screw attachments are affixed to each screw, one above and one below the ridge. The “bottom” of each screw attachment should contact the ridge. The screw attachments are held together with gel cyanoacrylate (“super glue”). Once affixed, the screw attachments should not move in the longitudinal axis of the screw, but should rotate freely with minimal resistance.
1.3) Assemble the micro-drive body (**Figure 2C-D**)
  1.3.1) Using fine, sharp scissors, cut “large” polyimide tubing (outer diameter 0.2921 mm, inner diameter 0.1803 mm) into approximately 6-cm-long sections.
  1.3.2) Pass the large polyimide sections through the output holes on the micro-drive cannula so that each tube extends beyond the “bottom” of the cannula by a few millimeters.
  1.3.3) Using a clean 30-gauge needle, affix the polyimide to the cannula by applying small amounts of liquid cyanoacrylate. Take care not to allow the cyanoacrylate to enter the inside of the polyimide tube. “Dripping” liquid cyanoacrylate down into the cannula through the top of the drive body can expedite this process, but will require re-clearing of the guide holes later with a fine-tipped drill.
  1.3.4) Pass the large polyimide tubes from the “top” of the micro-drive cannula through the appropriate large polyimide holes in the micro-drive body.
  1.3.5) Slowly push the micro-drive cannula and micro-drive body together until they are adjacent and the cannula/body attachment tabs interlock. Be careful not to kink or damage the polyimide tubes in the process. Each polyimide tube should smoothly pass from the bottom of the cannula out through the top of the micro-drive body. Slight bending is normal, but excessive bending of the polyimide tube can warp the tetrode and prevent it from passing straight into the brain.
  1.3.6) Affix the micro-drive body and the micro-drive cannula together using cyanoacrylate.
  1.3.7) Using a new, sharp razor blade, sever the large polyimide that are extruding from the “bottom” of the cannula output holes. The cut should be exactly at the base of the cannula, making the tubes and the cannula bottom flush with one another.
  1.3.8) Using sharp scissors, cut the large polyimide tubing just above the edge of the inner rim of the drive body at an approximately 45° angle.
1.4) Load the assembled custom screws (**Figure 2E**)
  1.4.1) Screw each assembled custom screw into the outer holes of the micro-drive body. Ensure that the screw guidepost passes through the large hole in the screw attachments. Advance each screw fully until it will not advance further. Pre-lubrication of screws with mineral oil or axle grease is recommended.
  1.4.2) Using extremely sharp scissors, cut “small” polyimide tubing (outer diameter 0.1397 mm, inner diameter 0.1016) into approximately 4-cm-long sections.
  1.4.3) Pass the small polyimide sections through the large polyimide already mounted in the micro-drive. There should be excess small polyimide protruding from both the top and bottom of each large polyimide tube.
  1.4.4) Affix the small polyimide tubes to the screw attachments via cyanoacrylate, taking care not to get any cyanoacrylate into the inside of either the large or small polyimide tube.
  1.4.5) Using a new, sharp razor blade, sever the small polyimide tube ends that are extruding from the “bottom” of the cannula holes. The cut should be exactly at the base of the cannula and the cut should be clean, with nothing blocking the polyimide tube hole.
  1.4.6) Using sharp scissors, cut the “top” of the small polyimide a few millimeters above the top of the screw attachment at an approximately 45° angle. The cut should be clean, with nothing blocking the polyimide tube hole.
1.5) Load the tetrodes
  1.5.1) Prepare tetrodes (~6 cm in length) using previously described methods^31^.
  1.5.2) Using ceramic- or rubber-tipped forceps, carefully pass a tetrode through one of the small polyimide tubes, leaving ~2 cm protruding from the “top” of the small polyimide tube.
  1.5.3) Affix the tetrode to the “top” of the small polyimide tube via liquid cyanoacrylate, taking care not to affix the small and large polyimide tubes together in the process.
  1.5.4) Retract the screw until it is near the top of the drive.
  1.5.5) Grab the tetrode wire protruding from the “bottom” of the drive and gently kink it at the point it is emerging from the cannula.
  1.5.6) Advance the screw fully back into the drive.
  1.5.7) Using very sharp scissors, cut the tetrode wire just above the kink. Under a microscope, ensure that the cut was clean and that the metal of all four tetrodes is exposed.
  1.5.8) Retract the screw until the tetrode is just secured within the cannula.
  1.5.9) Repeat 1.5.2 through 1.5.8 for all screws.
  1.5.10) Attach the EIB to the EIB support platform via small jewelry screws.
  1.5.11) Connect each electrode of each tetrode to the appropriate port on the EIB.
1.6) Prepare the micro-drive for surgery
  1.6.1) Electrically plate the tetrodes to reduce electrical impedence using previously described methods^31^.
  1.6.2) After plating, ensure that each tetrode is housed in the cannula such that the tip of the tetrode is flush with the “bottom” of each cannula hole.
  1.6.3) Slide the micro-drive cone around the completed micro-drive. Attach the micro-drive lid to the micro-drive cone by sliding the cone attachment pole into the lid port. Orient the cone so that the EIB connectors pass freely through the EIB connection pass-thru holes when the lid is closed and glue the cone in place with cyanoacrylate placed around the base of the cone, taking care not to get any cyanoacrylate into any of the cannula output holes. Remove the lid.
  1.6.4) Carefully backfill each cannula hole with sterile mineral oil to prevent bodily fluids from entering the polyimide holes after surgical implantation.
  1.6.5) Carefully coat the base of the cannula with sterile petroleum jelly. This will serve as a barrier to prevent chemical agents (*e.g.,* dental cement) from entering the exposed brain during the surgery.
  1.6.6) Weigh the fully assembled micro-drive, lid, and four bone screws to prepare an equal-weight counter-balance.
  1.6.7) Optionally, prior to the surgery, extrude the tetrodes a distance that is appropriate for reaching the target brain regions once the drive will be flush with the skull.

**Figure 1:**
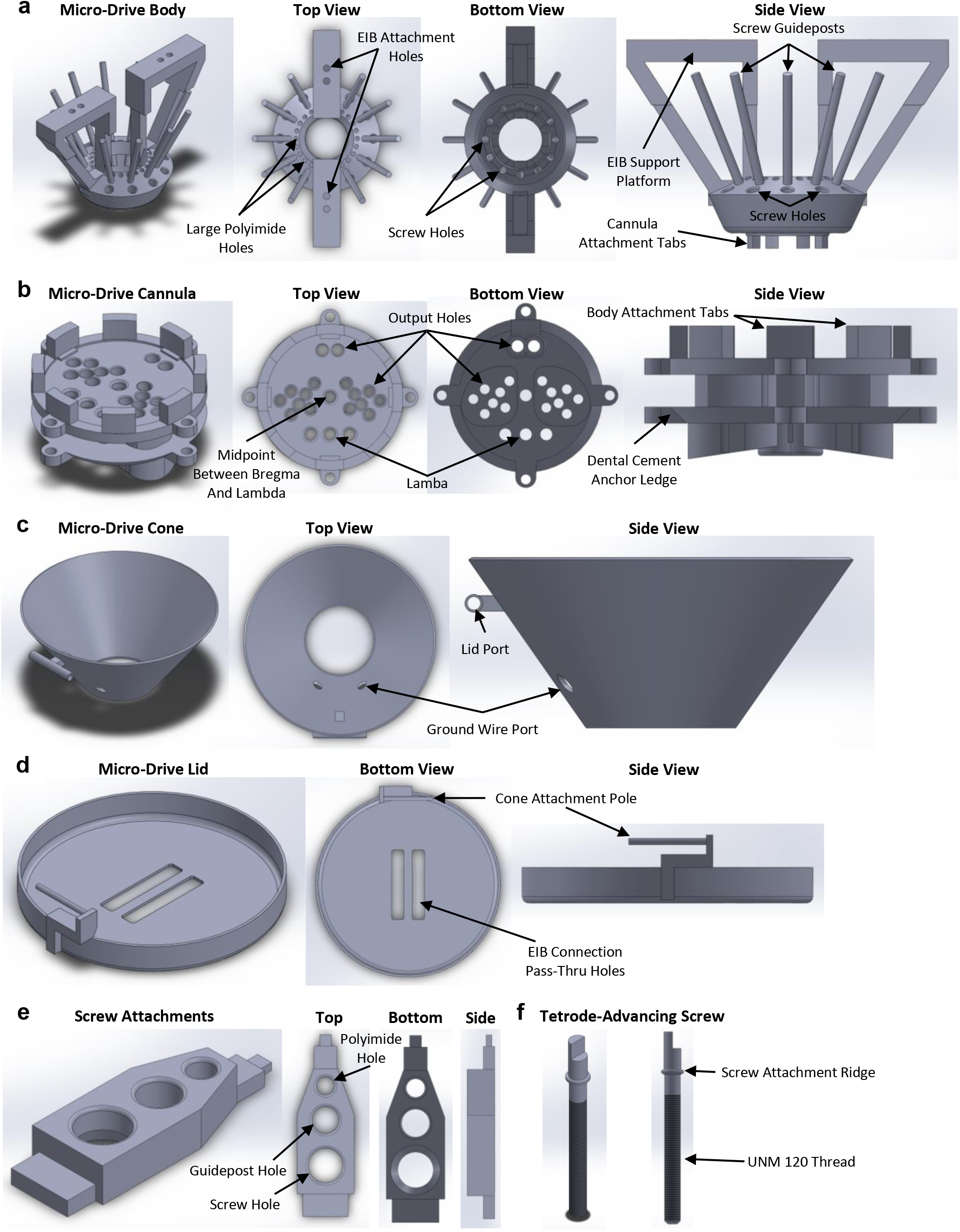
Micro-drive components. Three-dimensional renderings of micro-drive body (**A**), cannula (**B**), cone (**C**), lid (**D**), screw attachments (**E**), and tetrode-advancing screw (**F**). Critical features of each component are indicated. Measurement details can be extracted from the model files available at https://github.com/Brad-E-Pfeiffer/JuvenileMouseMicroDrive/releases/tag/v1.0.1.

**Figure 2:**
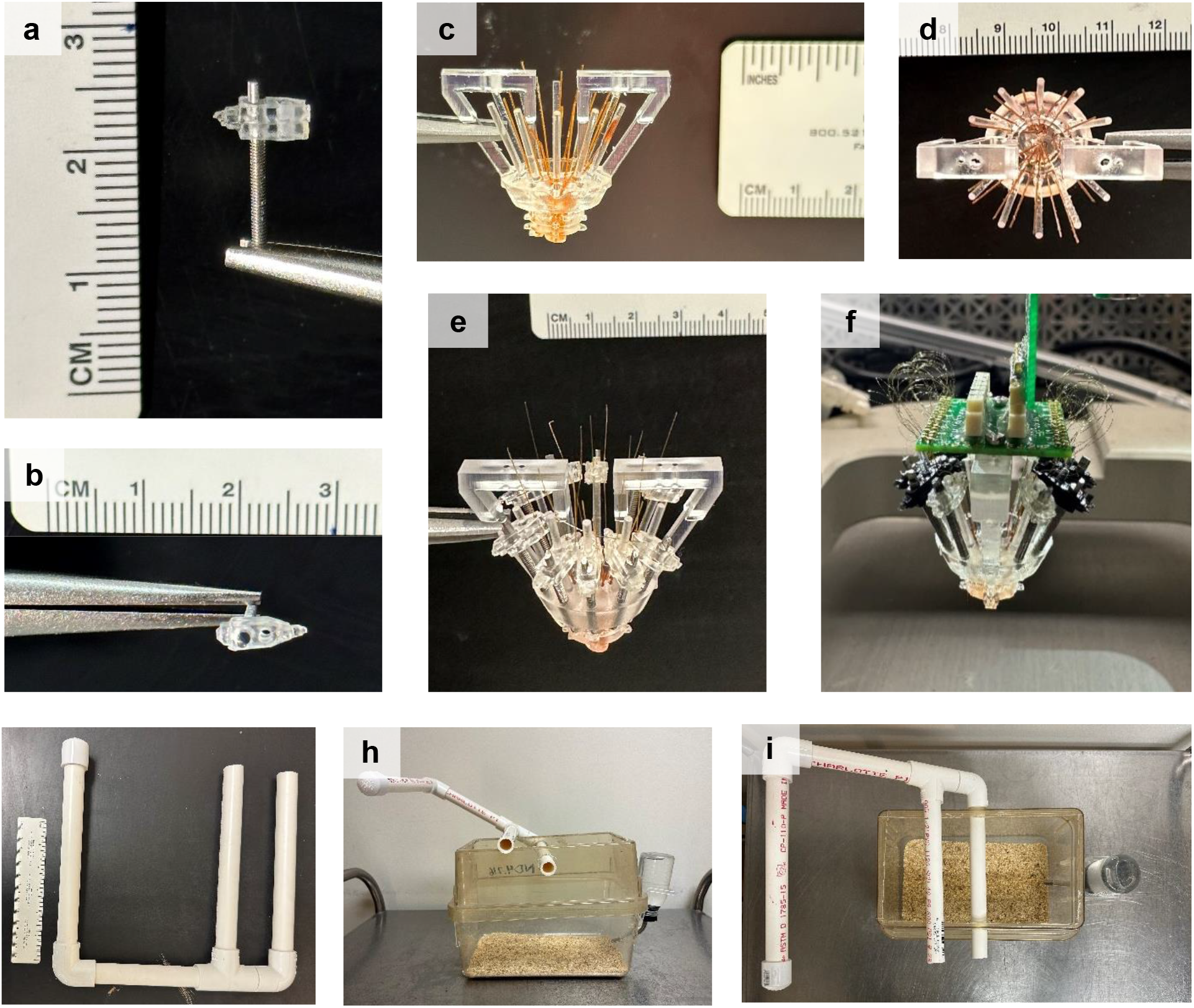
Micro-drive construction. **A-B.** Side (**A**) and top (**B**) view of tetrode-advancing screw with top and bottom screw attachments connected. **C-D.** Side (**C**) and top (**D**) view of micro-drive with body and cannula attached and large polyimide running through each cannula hole and trimmed to the bottom of the cannula. **E.** Side view of micro-drive with screws and small polyimide in place. The tops of the small polyimide tubes will be trimmed immediately prior to tetrode loading. **F**. Completed micro-drive attached to the stereotaxic apparatus. The protective cone that would normally surround the micro-drive has been removed for visualization purposes. Note that some of the screw attachments were printed in a black resin for this micro-drive. **G.** Counter-balance support system. **H-I.** Side (**H**) and top (**I**) view of mouse cage with counter-balance support system attached.

### 2. Surgical Implantation

2.1) Anesthetize the mouse and mount it in the stereotaxic apparatus
  2.1.1) Place mouse in a small box with sufficient space to move and anaesthetize the mouse with isoflurane. Other anesthetic agents can be used, but caution should be used due to the age, size, and weight of the juvenile mouse subject.
  2.1.2) Once the mouse is unresponsive (no response to tail pinch, ventilation rate approximately 60 breaths per minute), remove it from the box and quickly mount it on the stereotaxic apparatus.
  2.1.3) Quickly, place the stereotaxic anesthesia mask over the mouse’s snout and allow isoflurane to continue to maintain the subject in a fully anesthetized state.
  2.1.4) Fully secure the subject’s head in the stereotaxic apparatus using ear bars. Ensure the skull is level and immobile without placing unnecessary pressure to the subject’s ear canals. Due to limited ossification of juvenile skull bones, it is possible to cause permanent damage during head fixation.
2.2) Prepare the mouse for surgery and expose the skull
  2.2.1) Protect subject’s eyes by placing a small volume of synthetic tear gel on each eye and covering each eye with an autoclaved patch of foil. The synthetic tears will keep the eyes moist while the foil will prevent any light source from causing long-term damage. Thicker synthetic tear solutions are preferred as they can also serve as a barrier for inadvertent introduction of other potentially toxic surgical solutions (ethanol, dental acrylic, etc.) into the eyes.
  2.2.2) Remove the hair from the scalp using hair-removal cream, being exceedingly careful not to get the gel near the eyes. Use sterile cotton-tipped swabs to apply the gel and remove the gel and hair.
  2.2.3) Using sterile cotton-tipped swabs, clean the scalp via three consecutive washes of povidone-iodine (10%) solution followed by isopropyl alcohol (100%).
  2.2.4) Using a sterile scalpel or fine scissors, remove the scalp.
  2.2.5) Using sterile cotton-tipped swabs and sterile solutions of saline solution (0.9% NaCl) and hydrogen peroxide, thoroughly clean the skull.
  2.2.6) Identify bregma and, using the stereotaxic apparatus, carefully mark target recording locations on the skull with a permanent marker.
2.3) Open cannula hole and attach bone anchors
  2.3.1) Remove the skull overlaying the recording sites. Due to the thinness of the skull at this age, the skull can be cut with a scalpel blade, removing the necessity of a drill that may damage the underlying dura. Keep the exposed dura moist with application of sterile saline solution (0.9% NaCl) or sterile mineral oil. The dura does not need to be removed or punctured at this stage, as it is sufficiently thin in juvenile mice for the tetrodes to pass through in future steps.
  2.3.2) Carefully drill pilot holes for four bone screws. Bone screw placement should be in the extreme lateral and rostral or caudal portions of the skull, where the bone is thickest and the bone screw is sufficiently far away from the micro-drive implant. Bone screws should be sterile, fine jewelry screws (*e.g.,* UNM 120 thread, 1.5-mm head).
  2.3.3) Using a scalpel blade or carefully with a drill bit, score the skull near the bone screw hole locations. Scoring is important to provide a sufficiently rough surface for the liquid cyanoacrylate to bind in 2.3.5.
  2.3.4) Thread each bone screw into place, taking care not to pierce the underlying dura.
  2.3.5) Using a sterile 30-gauge needle, place liquid cyanoacrylate around each bone screw. This effectively thickens the skull where the bone screws have been attached. Take care not to get any cyanoacrylate into the exposed dura above the recording sites.
2.4) Lower and attach micro-drive (**Figure 2G**)
  2.4.1) Mount the completed micro-drive onto the stereotaxic apparatus so that it can be carefully lowered onto the mouse skull. Ensure that when lowered, the micro-drive cannula will be at the appropriate coordinates.
  2.4.2) Lower the micro-drive slowly, moving only in the dorsal/ventral direction. We prefer to lower the micro-drive with the tetrodes already advanced out of the cannula holes (step 1.6.5) in order to visualize their entry into the brain, and any medial/lateral or rostral/caudal movement when the tetrodes are touching the mouse can bend the tetrodes and cause them to miss their final destination.
  2.4.3) Once the micro-drive is fully lowered, the base of the cannula should just make contact with the skull/dura. The layer of petroleum jelly and/or mineral oil will serve as a barrier to cover the exposed dura. If necessary, ensure that excess exposed dura is covered by adding additional sterile petroleum jelly or sterile bone wax.
  2.4.4) While holding the micro-drive in place with the stereotaxic apparatus, coat the skull in dental cement to affix the micro-drive base to the implanted bone screws. Dental cement should fully encase all bone screws and should cover the dental cement anchor ledge on the micro-drive cannula.
  2.4.5) While the dental cement dries, carefully shape it to prevent sharp corners or edges that may harm the mouse or damage the micro-drive. Ensure there is sufficient dental cement to hold the micro-drive, but eliminate excessive dental cement that will add unnecessary weight.
  2.4.6) Once the dental cement is fully set, carefully detach the micro-drive from the stereotaxic apparatus. Place the lid on the micro-drive.
  2.4.7) With a sterile cotton-tipped swab and sterile saline, clean the mouse.
  2.4.8) With a sterile cotton-tipped swab, apply a thin layer of anti-biotic ointment to any exposed scalp near the implant site.
  2.4.9) Remove the foil from the mouse’s eyes.
  2.4.10) Remove the mouse from the stereotaxic apparatus, taking care to support the additional weight of the micro-drive as the mouse is transported to a clean cage.

### 3. Post-Surgery Recovery

3.1) Immediate recovery
  3.1.1) Prior to the surgery, prepare the counter-balance system by connecting 0.75-inch-diameter PVC pipe as shown in **Figure 2G**. One arm of the system passes through holes drilled into the lid of the cage, the second arm rests on top of the cage lid, and the third arm extends above and beyond the cage. The top-most arm is capped.
  3.1.2) Before the mouse recovers from the surgery anesthesia, apply any veterinary-approved pain relief or anti-inflammatory agents.
  3.1.3) Carefully attach the micro-drive to the counter-balance system (**Figure 2G-I**), using a counter-balance weight identical to the weight of the micro-drive and bone screws. A fishing line runs from a connector attached to the EIB, over the three arms of the counter-balance system, to the counter-balance weight which hangs over the top-most arm.
  3.1.4) Ensure that the counter-balance is strongly connected to the micro-drive EIB and that there is sufficient line to give the mouse complete access to the entirety of the cage.
  3.1.5) Provide nutrient-rich gel in the cage alongside moistened normal rodent chow to ensure rehydration and recovery.
  3.1.6) Monitor the mouse until it fully recovers from the surgery anesthesia.
3.2) Long-term recovery
  3.2.1) At all times when not attached to the recording equipment, the micro-drive should be supported by the counter-balance system. The counter-weight can be reduced over time, but should never be removed entirely to avoid unanticipated stress to the mouse or torque to the bone screws.
  3.2.2) Provide nutrient-rich gel for at least three days following surgery, at which point solid food alone will be sufficient. Because of the overhead requirements of the counter-balance system, food and water cannot be provided in an overhead wire mesh and food must be placed on the cage floor and water must be provided through the side of the cage. To prevent spoilage, food must be entirely replaced daily.
  3.2.3) Daily, ensure that the mouse has free access to the entirety of the cage, with counter-balance robustly and strongly attached to the micro-drive.

## REPRESENTATIVE RESULTS

We have used the above-described protocol to successfully record local field potential signals and single units from multiple brain areas simultaneously in mice, with daily recordings from the same mice from p20 to p60. Here, we report representative electrophysiological recordings from two mice and post-experiment histology demonstrating final recording locations.

### Surgical Implantation of Micro-drive into p20 mice

A micro-drive (**Figure 1**) was constructed (**Figure 2**) and surgically implanted into a p20 mouse as described above. Immediately following the surgery, the mouse was attached to the counter-balance system (**Figure 2G-I**) and allowed to recover. Once the mouse was fully mobile, the micro-drive was plugged into a Neuralynx recording system. The weight of the cables connecting the micro-drive to the recording equipment was suspended above the mouse. Electrophysiological recordings (32 kHz) were obtained across all channels for one hour while the mouse behaved naturally in its home cage. Following recording, the mouse was unplugged from the recording system, reattached to the counter-balance system, and returned to the vivarium with free access to water and chow.

### Daily recording of neural activity

Electrophysiological recordings were obtained daily for several weeks, providing chronic monitoring of the same brain region across the critical developmental windows of p20-p60. Sample raw local field potentials (LFP) from across the chronic recordings are shown in **Figure 3A,C**. Isolated single units were simultaneously obtained from multiple tetrodes (**Figure 3B**). Units with similar waveforms were identified across multiple days (**Figure 3B, middle and right)**, but due to possible drift of the recording electrode, we are hesitant to claim that the same unit is being identified across days. These data demonstrate stable, high-quality *in vivo* electrophysiological recordings from the same mouse across early development.

**Figure 3:**
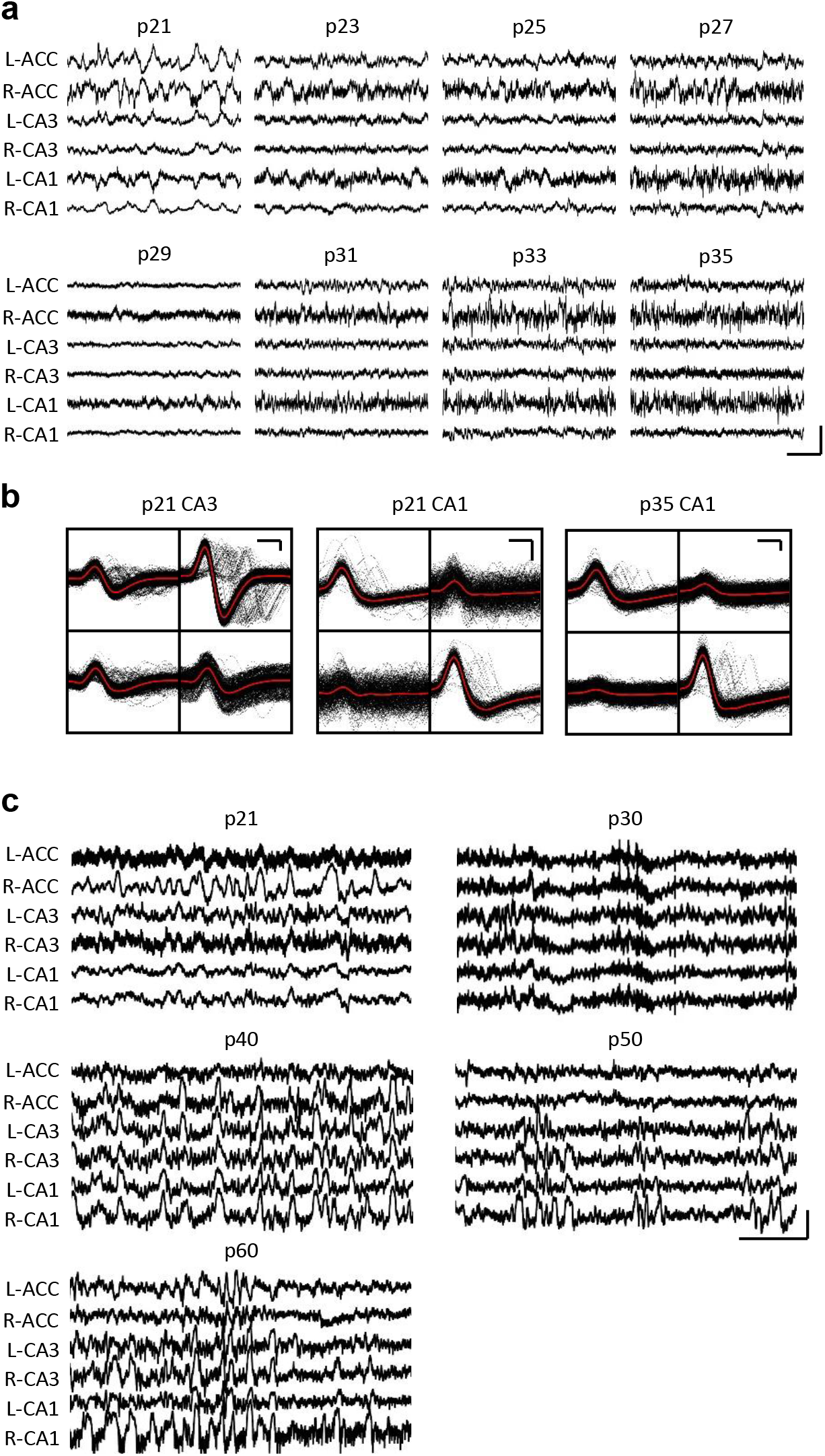
Representative electrophysiological recordings. A p20 mouse was implanted with a micro-drive as described above. Starting on p21 and every day thereafter for two weeks, the mouse was attached to the recording apparatus and neural activity was recorded for at least one hour. **A.** Raw local field potential (LFP) recordings from bilateral (L = Left; R = Right) anterior cingulate cortex (ACC), hippocampal area CA3 (CA3) and hippocampal area CA1 (CA1). Data were collected every day; for clarity, only data from odd days are displayed. All traces are taking during periods of immobility in the home cage. Scale bar 1 mV, 2 s. **B.** For the recordings in **A**, representative single units isolated from hippocampal area CA3 (left) and CA1 (right). All raw waveforms on each electrode are shown in black; average in red. Scale bar 50 μV, 0.2 ms. **C.** For a second mouse implanted at p20, representative raw LFP traces for every tenth day until final recording day at p60. Data were collected every day; for clarity, only data from every tenth day are displayed. All traces are taking during periods of immobility in the home cage. Scale bar 1 mV, 2 s.

### Histological confirmation of recording sites

Following the final recording day, the mouse was thoroughly anesthetized and current was passed through the electrode tips to produce small lesions at the recording sites. Post-experiment histological sectioning of the mouse brain allowed visualization of the final recording sites (**Figure 4A-B**).

**Figure 4:**
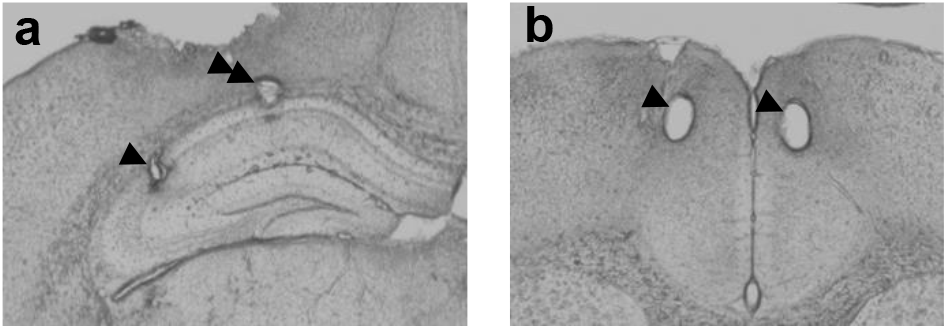
Representative histology. A p20 mouse was implanted with a micro-drive as described above. Following the final recording day on p60, electrolytic lesions were produced at the recording sites and the brain was perfused with 4% paraformaldehyde. 50 μm sections were produced to identify the recording sites. **A.** Lesions in CA1 and CA3 of hippocampus Arrowhead denotes CA3 recording site; double-arrowhead denotes CA1 recording site. **B.** Lesions in bilateral ACC. Arrowheads denote ACC recording sites.

## DISCUSSION

Modern experiments exploring *in vivo* neural circuit function in rodents often utilize extracellular electrophysiology via permanently implanted electrodes to monitor the activity of individual neurons (*i.e.,* single units) or local populations (via local field potentials, LFP), but such methods are rarely applied to juvenile mice due to technical challenges. Here, we describe a novel method for obtaining *in vivo* electrophysiological recordings in mice across developmentally critical windows of p20 to p60 and beyond. Our methodology involves a novel manufacturing process for printing and construction of a micro-drive implant, a surgical implantation procedure, and a post-surgery recovery strategy, all specifically tailored for use in juvenile mice. Several considerations were influential in our protocol, including the small size and relative weakness of juvenile mice compared to their adult counterparts as well as the reduced ossification of the juvenile mouse skull on which the micro-drive needed to be attached.

Two primary methods commonly used to perform *in vivo* electrophysiology are arrays of electrodes (*e.g.*, tetrodes) and silicon probes. Silicon probes are lightweight, can provide a large number of recording sites per unit weight, and have been previously utilized in juvenile rats^25^. However, silicon probes are relatively expensive per unit. In contrast, our micro-drive can be constructed using less than $50 USD in raw materials, making it a cost-effective option for *in vivo* recording. In addition, silicon probes must often be implanted in fixed lines, prohibiting recordings of spatially diverse brains regions. In contrast, our micro-drive design utilizes independently adjustable tetrodes to accommodate simultaneous recordings in up to sixteen different locations with virtually no restriction on the spatial relationship between those locations. Our micro-drive design can be easily modified to support targeting of different locations than those described here by moving the cannula hole extrusions to any desired anterior/posterior and medial/distal location. When targeting alternate brain areas, it is important to note that while tetrodes will often travel straight, it is possible for these thin wires to deflect slightly as they exit the micro-drive cannula. Thus, the smaller or more ventral a brain region is, the more challenging it will be to successfully target the area with tetrodes.

To maintain a consistent recording across multiple days, wires or probes must be rigidly affixed to the skull. While the overall structure of the mouse skull undergoes only minor changes after p20, the skull thickens considerably between ages p20 and p45 ^32^. Indeed, the skull at p20 is insufficiently rigid to support an attached implant without being damaged. To overcome this biological limitation, we artificially thicken the skull via cyanoacrylate. Implantation in mice younger than p20 is likely possible using this strategy, but the mouse skull displays considerable size and shape changes until roughly p20^32^. Thus, we do not recommend implantation for extended periods in mice younger than p20 as the cyanoacrylate and fixed bone screws in the still-developing skull may significantly impact the natural growth of the skull and underlying brain tissue development.

A critical step in our method is the post-surgery recovery strategy, in which the weight of the implant is persistently counter-balanced as the mouse matures and undergoes muscular and musculoskeletal system development. Early after implantation, mice are unable to successfully bear the weight of the implant without the counter-balance, leading to malnourishment and dehydration as the mouse cannot adequately reach food and water sources in its cage. Our counter-balance system is easy and inexpensive to construct, trivial to implement, and allows mice of any implantable age to freely explore the entirety of their home cage, ensuring adequate nutrition and hydration. As mice age, the amount of counter-balance can be lessened until it can be entirely removed in adult mice; however, we recommend continued use of the counter-balance system for the duration of the experiment with at least a nominal counterweight attached at all times. While an adult mouse may be able to bear the size and weight of the micro-drive over time, continued natural movement during free behavior with no ameliorating counterweight produces torque and shearing force on the bone screws that anchor the micro-drive onto the skull, making it increasingly likely to become detached, especially during longer chronic experiments.

Many human neurological and psychiatric disorders manifest during periods of early development or across adolescence, including autism and schizophrenia. However, we know little regarding the circuit-level dysfunction which may underlie these diseases, despite a plethora of mouse models. Identification of these initial network changes is critical for creating early detection strategies and treatment paradigms. Yet, due to technical challenges, it remains unclear how network function is disrupted across development in mouse models of neuro-psychiatric diseases. The novel micro-drive and recovery strategy we describe here is designed to support investigations into multi-regional brain network development in the mouse brain, allowing researchers to measure healthy brain development as well as identify alterations to that development in mouse models of disease.

## ACKNOWLEDGMENTS

This work was supported by National Institutes of Health R01 NS104829 (BEP), R01 MH117149 (LJV), F99NS12053 (LDQ), and UT Southwestern GSO Endowment Award (RJP and LDQ). We thank Jenny Scaria (Texas Tech University Health Sciences Center School of Pharmacy) for technical assistance and Dr. Brendon Watson (University of Michigan) for methodological suggestions.

## DISCLOSURES

The authors have nothing to disclose.

## Notes

### Competing Interest Statement

The authors have declared no competing interest.

## REFERENCES

1. Konrad, K., Firk, C., Uhlhaas, P.J. Brain development during adolescence. Deutsches Arzteblatt International. 110 (25), 425–431, doi: 10.3238/arztebl.2013.0425 (2013).

2. Silbereis, J.C., Pochareddy, S., Zhu, Y., Li, M., Sestan, N. The cellular and molecular landscapes of the developing human central nervous system. Neuron. 89 (2), 248–268, doi: 10.1016/j.neuron.2015.12.008 (2016).

3. Semple, B.D., Blomgren, K., Gimlin, K., Ferriero, D.M., Noble-Haeusslein, L.J. Brain development in rodents and humans: Identifying benchmarks of maturation and vulnerability to injury across species. Progress in Neurobiology. 106–107, 1–16, doi: 10.1016/j.pneurobio.2013.04.001 (2013).

4. Volk, L., Chiu, S.-L., Sharma, K., Huganir, R.L. Glutamate synapses in human cognitive disorders. Annual Review of Neuroscience. 38, 127–149, doi: 10.1146/annurev-neuro-071714-033821 (2015).

5. Lord, C. et al. Autism spectrum disorder. Nature Reviews Disease Primers. 6 (1), 1–23, doi: 10.1038/s41572-019-0138-4 (2020).

6. McCutcheon, R.A., Reis Marques, T., Howes, O.D. Schizophrenia - An overview. JAMA Psychiatry. 77 (2), 201–210, doi: 10.1001/jamapsychiatry.2019.3360 (2020).

7. Hopfield, J.J. Neural networks and physical systems with emergent collective computational abilities. Proceedings of the National Academy of Sciences. 79 (8), 2554–2558, doi: 10.1073/pnas.79.8.2554 (1982).

8. Heeger, D.J. Theory of cortical function. Proceedings of the National Academy of Sciences. 114 (8), 1773–1782, doi: 10.1073/pnas.1619788114 (2017).

9. Pouget, A., Dayan, P., Zemel, R. Information processing with population codes. Nature Reviews Neuroscience. 1, 125–132 (2000).

10. Averbeck, B.B., Latham, P.E., Pouget, A. Neural correlations, population coding and computation. Nature Reviews Neuroscience. 7 (5), 358–366, doi: 10.1038/nrn1888 (2006).

11. Bey, A.L., Jiang, Y. hui Overview of mouse models of autism spectrum disorders. Current Protocols in Pharmacology. 2014(September), 5.66.1–5.66.26, doi: 10.1002/0471141755.ph0566s66 (2014).

12. Kazdoba, T.M., Leach, P.T., Yang, M., Silverman, J.L., Solomon, M., Crawley, J.N. Translational mouse models of autism: Advancing toward pharmacological therapeutics. Currrent Topics in Behavioral Neurosciences. 28, 1–52, doi: 10.1007/7854 (2016).

13. Mendoza, M.L., Quigley, L.D., Dunham, T., Volk, L.J. KIBRA regulates activity-induced AMPA receptor expression and synaptic plasticity in an age-dependent manner. iScience. 25 (12), 105623, doi: 10.1016/j.isci.2022.105623 (2022).

14. Bernardet, M., Crusio, W.E. Fmr1 KO mice as a possible model of autistic features. The Scientific World Journal.6, 1164–1176, doi: 10.1100/tsw.2006.220 (2006).

15. Weaving, L.S., Ellaway, C.J., Gécz, J., Christodoulou, J. Rett syndrome: Clinical review and genetic update. Journal of Medical Genetics. 42 (1), 1–7, doi: 10.1136/jmg.2004.027730 (2005).

16. Krawczyk, M. et al. Hippocampal hyperexcitability in fetal alcohol spectrum disorder: Pathological sharp waves and excitatory/inhibitory synaptic imbalance. Experimental Neurology.280, 70–79, doi: 10.1016/j.expneurol.2016.03.013 (2016).

17. Jaramillo, T.C., Speed, H.E., Xuan, Z., Reimers, J.M., Liu, S., Powell, C.M. Altered striatal synaptic function and abnormal behaviour in Shank3 exon4-9 deletion mouse model of autism. Autism Research. 9 (3), 350–375, doi: 10.1002/aur.1529 (2016).

18. Suh, J., Foster, D.J., Davoudi, H., Wilson, M.A., Tonegawa, S. Impaired hippocampal ripple-associated replay in a mouse model of schizophrenia. Neuron. 80 (2), 484–493, doi: 10.1016/j.neuron.2013.09.014 (2013).

19. Altimus, C., Harrold, J., Jaaro-Peled, H., Sawa, A., Foster, D.J. Disordered ripples are a common feature of genetically distinct mouse models relevant to schizophrenia. Molecular Neuropsychiatry. 1, 52–59, doi: 10.1159/000380765 (2015).

20. Marcotte, E.R., Pearson, D.M., Srivastava, L.K. Animal models of schizophrenia: A critical review. Journal of Psychiatry and Neuroscience. 26 (5), 395–410 (2001).

21. Makuch, L. et al. Regulation of AMPA receptor function by the human memory-associated gene KIBRA. Neuron. 71 (6), 1022–1029, doi: 10.1016/j.neuron.2011.08.017 (2011).

22. Dutta, S., Sengupta, P. Men and mice: Relating their ages. Life Sciences.152, 244–248, doi: 10.1016/j.lfs.2015.10.025 (2016).

23. Kloosterman, F. et al. Micro-drive array for chronic in vivo recording: Drive fabrication. Journal of Visualized Experiments. (26), 1–4, doi: 10.3791/1094 (2009).

24. Buzsáki, G. Large-scale recording of neuronal ensembles. Nature Neuroscience. 7 (5), 446–51, doi: 10.1038/nn1233 (2004).

25. Farooq, U., Dragoi, G. Emergence of preconfigured and plastic time-compressed sequences in early postnatal development. Science. 363 (6423), 168–173, doi: 10.1126/science.aav0502 (2019).

26. Langston, R.F. et al. Development of the spatial representation system in the rat. Science. 328 (5985), 1576–1580, doi: 10.1016/b978-0-408-01434-2.50020-6 (2010).

27. Wills, T.J., Cacucci, F., Burgess, N., O’Keefe, J. Development of the hippocampal cognitive map in preweanling rats. Science. 328 (5985), 1573–1576, doi: 10.1126/science.1188224 (2010).

28. Bjerknes, T.L., Moser, E.I., Moser, M.B. Representation of geometric borders in the developing rat. Neuron. 82 (1), 71–78, doi: 10.1016/j.neuron.2014.02.014 (2014).

29. Bjerknes, T.L., Dagslott, N.C., Moser, E.I., Moser, M.-B. Path integration in place cells of developing rats. Proceedings of the National Academy of Sciences. 115 (7), E1637–E1646, doi: 10.1073/pnas.1719054115 (2018).

30. Jansen, N.A. et al. Impaired θ-γ coupling indicates inhibitory dysfunction and seizure risk in a Dravet syndrome mouse model. Journal of Neuroscience. 41 (3), 524–537, doi: 10.1523/JNEUROSCI.2132-20.2020 (2021).

31. Nguyen, D.P. et al. Micro-drive array for chronic in vivo recording: Tetrode assembly. Journal of Visualized Experiments. (26), 7–9, doi: 10.3791/1098 (2009).

32. Vora, S.R., Camci, E.D., Cox, T.C. Postnatal ontogeny of the cranial base and craniofacial skeleton in male C57BL/6J mice: A reference standard for quantitative analysis. Frontiers in Physiology.6(JAN), 1–14, doi: 10.3389/fphys.2015.00417 (2016).

